# Application of rapid invisible frequency tagging for brain computer interfaces

**DOI:** 10.1101/2022.05.09.491156

**Authors:** Marion Brickwedde, Yulia Bezsudnova, Anna Kowalczyk, Ole Jensen, Alexander Zhigalov

## Abstract

**Background:** Brain-computer interfaces (BCI) based on steady-state visual evoked potentials (SSVEPs/SSVEFs) are among the most commonly used BCI systems. They require participants to covertly attend to visual objects flickering at specified frequencies. The attended location is decoded in real-time by analysing the power of neuronal responses at the flicker frequency.

**New method:** We implemented a novel rapid invisible frequency-tagging technique, utilizing a state-of-the-art projector with refresh rates of up to 1440 Hz. We flickered the luminance of visual objects at 56 and 60 Hz, which was invisible to participants but produced strong neuronal responses measurable with magnetoencephalography (MEG). The direction of covert attention, decoded from frequency-tagging responses, was used to control a real-time BCI PONG game.

**Results:** Our results show that seven out of eight participants were able to play the pong game controlled by the frequency-tagging signal, with average accuracies exceeding 60%. Importantly, participants were able to modulate the power of the frequency-tagging response within a 1-second interval, while only seven occipital sensors were required to reliably decode the neuronal response.

**Comparison with existing methods:** In contrast to existing SSVEP-based BCI systems, rapid frequency-tagging does not produce a visible flicker. This extends the time-period participants can use it without fatigue, by avoiding distracting visual input. Furthermore, higher frequencies increase the temporal resolution of decoding, resulting in higher communication rates.

**Conclusion:** Using rapid invisible frequency-tagging opens new avenues for fundamental research and practical applications. In combination with novel optically pumped magnetometers (OPMs), it could facilitate the development of high-speed and mobile next-generation BCI systems.

## 1. Introduction

In recent years, brain-computer interfaces (BCIs) have developed from niche applications to wide implementations in clinical and rehabilitation settings as well as fundamental research paradigms (Bagherzadeh et al., 2020; Bergmann et al., 2019; Biasiucci et al., 2018; Brickwedde et al., 2019; Peles et al., 2020; Zrenner et al., 2018). One of the most common types of BCIs is based on steady-state visual evoked potentials or fields (SSVEPs/SSVEFs, e.g., Cheng et al., 2002; Kelly et al., 2005; Ma et al., 2022). These are characterised by rhythmic neuronal activity at the frequency of sensory stimulation. Specifically, flicker or frequency-tagging at a specific frequency will evoke neuronal responses echoing the pattern of stimulation (Brickwedde et al., 2020; Colon et al., 2012; Müller & Hillyard, 2000; Regan, 1982; Snyder, 1992; Stapells et al., 1984; Zhigalov et al., 2019). The magnitude of SSVEPs/SSVEFs is known to be attention-dependent (Müller et al., 1998; Müller & Hillyard, 2000; Toffanin et al., 2009; Zhigalov et al., 2019), which can be utilized to provide human participants with control over BCI systems.

However, most of the SSVEP/SSVEF BCI systems utilise stimulation at relatively low frequencies, usually within a range of 5 – 40 Hz, making the flickering clearly visible to participants (Landis, 1954; Seitz et al., 2005). The resulting visual flicker can be very distracting, straining to the eye and cause fatigue. Another important consideration pertains to epileptic seizures, which are most strongly provoked by lower frequency flicker in the range of 15 – 25 Hz (Fisher et al., 2005).

While the magnitude of SSVEPs/SSVEFs responses decreases with an increase in visual stimulation frequency (Regan, 1977), the temporal resolution increases greatly by increasing the number of queries and therefore neuronal responses in a given temporal interval. Thus, the use of higher frequencies for visual stimulation could greatly enhance the communication rate compared to existing BCI systems.

Intrinsic oscillatory activity in lower frequency ranges is dominant across all sensory cortices and it is believed to fulfil crucial functional roles for sensory processing and memory (Haegens & Golumbic, 2018; Jensen & Mazaheri, 2010). As such, scalp-level measurements during low-frequency sensory stimulation can produce a signal mix of cortical oscillatory activity and SSVEPs/SSVEFs, which can become difficult to disentangle. At the same time, low-frequency SSVEP/SSVEF activity can affect and entrain intrinsic oscillatory activity (Keitel et al., 2014; Spaak et al., 2014), which can influence task performance and question the interpretability of fundamental research analyses concerning oscillatory activity.

State-of-the-art projectors are now able to produce refresh rates of up to 1440 Hz. This technological advancement has introduced the possibility to implement rapid invisible frequency tagging (RIFT), which is attention-dependent comparable to its low-frequency counterparts, but invisible to participants (Drijvers et al., 2021; Seijdel et al., 2022; Zhigalov et al., 2019; Zhigalov & Jensen, 2020). It does not entrain intrinsic oscillatory activity (Duecker et al., 2021) and elicits very localized responses (Zhigalov et al., 2019; Zhigalov & Jensen, 2020), diminishing potential confounds of task performance and interpretability of low-frequency activity, as well as introducing the possibility to work with a limited number of sensors.

Therefore, we implemented rapid-invisible frequency tagging into a real-time BCI system to provide proof-of-principle for the validity of this technique. Participants used covert attention while fixating a cross at the centre of the screen, to control a quickly paced pong game. We were interested in the overall performance of participants and the efficacy of our system to track visuo-spatial attention in real-time.

## 2. Materials and Methods

### 2.1. Participants

Eight healthy participants with previous experience in MEG studies were recruited at the University of Birmingham (5 women, mean age: 32 ± 3). Participants had normal or corrected-to-normal vision and no history of neurological disorders. The study protocol was approved by the Ethics Committee of the School of Psychology at Birmingham University and in accordance with the Declaration of Helsinki. All participants provided written informed consent.

### 2.2. Setup

Participants were seated comfortably in the gantry of a 306-sensor TRIUX Elekta system with 204 orthogonal planar gradiometers and 102 magnetometers (Elekta, Finland), in a dimly lit magnetically shielded room. We used a PROPixx DLP LED projector (VPixx Technologies Inc., Canada) to present visual stimuli during training and game sessions. Importantly, the projector supported a refresh rate of 1440 Hz, which allowed the application of high-frequency flicker or tagging, which was invisible to participants. The video output was projected onto a 71 × 40 cm screen placed 1.5 m away from the participant. During the experiment, horizontal and vertical eye movements were tracked with EyeLink 1000 Plus, SR Research Ltd, Canada. MEG signals passed through an embedded analogue filter, 0.1 – 330 Hz, and sampled at 1000 Hz. Different types of data, including MEG, eye-tracking, and triggers, were combined into a single data-stream in the acquisition computer. The combined data were then split into 100 ms blocks and sent to the stimulus computer using the Fieldtrip buffer (Oostenveld et al., 2011). Finally, these data were used to train a classifier to decode the direction of covert attention based on the power of the frequency-tagging response.

### 2.3. BCI-pong training

Before the training session, the eye-tracker was calibrated by asking participants to sequentially fixate on different spatial locations across the screen. This information was then used to automatically discard trials with eye-blinks during the frequency-tagging phase of each trial. During training, participants were asked to fixate on the cross at the centre of the screen. Each trial started with 1 s baseline period. Afterwards, a grey ball was displayed for 2 s at a random location on either the right or the left side of the screen. At the same time, the left and the right sides of the screen (see, Fig. 1, dashed line) were frequency-tagged at 56 and 60 Hz, respectively. The edges of the tagged patches were 100% transparent, slowly fading in to 100% visibility over 10% of the length/height of the patch. A static vertical bar of ∼2 degrees of visual angle was displayed in the lower part of the screen (Fig. 1), to make the training session as close to the subsequent gaming session as possible.

**Fig. 1.**
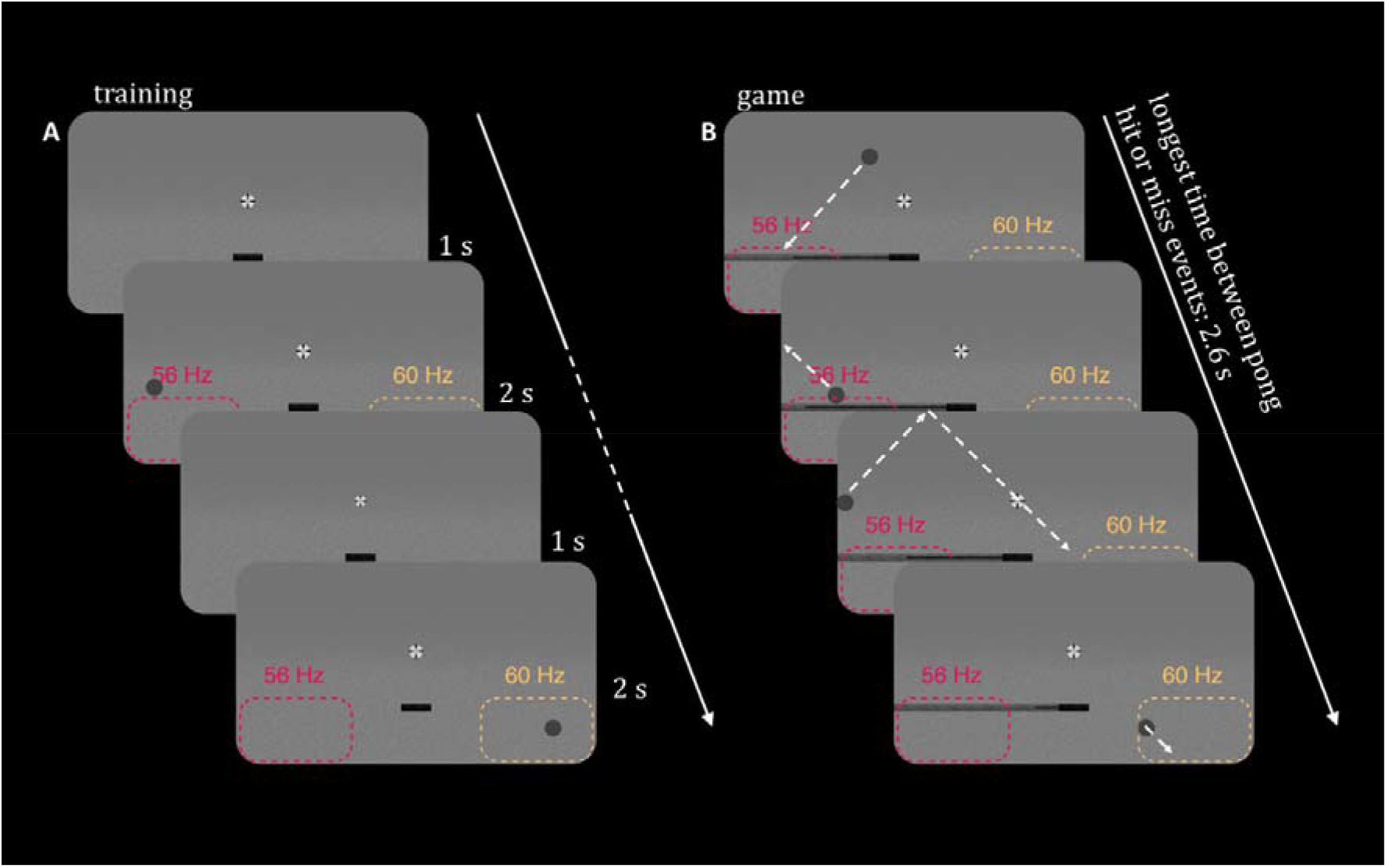
Experimental setup of the BCI-pong training and game. (A) In the training session, participants were asked to covertly attend either left or right side of the screen for 2 s. At the same time, invisible flickering patches were presented at the right and left bottom of the screen. Dashed areas indicate the location of flickering patches and corresponding tagging frequencies. (B) During the game, participants covertly attended the right or left side of the screen, to move a bar from one side of the screen to another. The goal was to always position the bar at the side of the screen the pong ball was falling towards. Inside of the bar, a dark line displayed the continuous classifier output as a quantitative measure of covert attention. Additionally, a small black bar in the middle was always present.

Participants were asked to keep fixating on the cross, while covertly attending to the side of the screen indicated by the location of the pong ball (see Fig. 1). Additionally, participants were asked to reduce blinking as much as possible and to limit blinking to the end of the trial. All participants performed 80 valid trials (40 to the left and right, randomly sequenced). The training was performed once at the beginning of the experiment.

MEG data acquired in the training session were used to train a classifier (see, Classification algorithm section).

### 2.4. BCI-pong game

In the game, participants controlled the position of the horizontal bar by changing the direction of their covert attention (Fig. 1). The goal of the game was to keep the pong ball afloat, by moving the bar to the right or the left side of the screen. During the game, real-time MEG data were stored in a 1-s buffer, which was updated every 100 ms. This way, we assessed the power spectral density of the frequency-tagging response using Fourier transform (1), as implemented in MATLAB

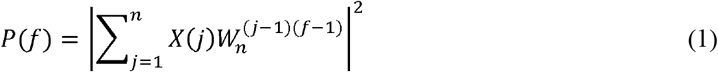

where

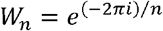

*P* is the power spectral density, *X* denotes single-sensor MEG signal, *W*_*n*_ is one of *n* roots of unity, *n* is the signal length.

The power of the tagged response at 56 and 60 Hz was used in the classifier, to infer the direction of covert attention in real-time. The processing time was negligibly small (less than 10 ms on average) compared to the data acquisition delay (100 ms) which ensured uninterrupted feedback for the user.

The game started with the pong ball appearing at a random position in the top quarter of the screen, moving towards the bottom. If the bar and the ball were at the same side of the screen and colliding, the ball was reflected back in a 90° angle and moved towards the wall. There, it bounced off until it was moving towards the bottom again. If the bar was on the opposite side when the ball reached collision height, the ball fell through the bottom and was reset to a random starting position at the top (see Fig. 2). The first second after a reset, the ball always moved with reduced speed to accommodate for an orientation period necessary to predict where the ball was moving towards. If participants blinked in the last second before the collision, the trial was discarded. Each participant played the 5-minute BCI-pong game four times. At the end of each game, a score indicating the number of hits and misses was presented on the screen.

**Fig. 2.**
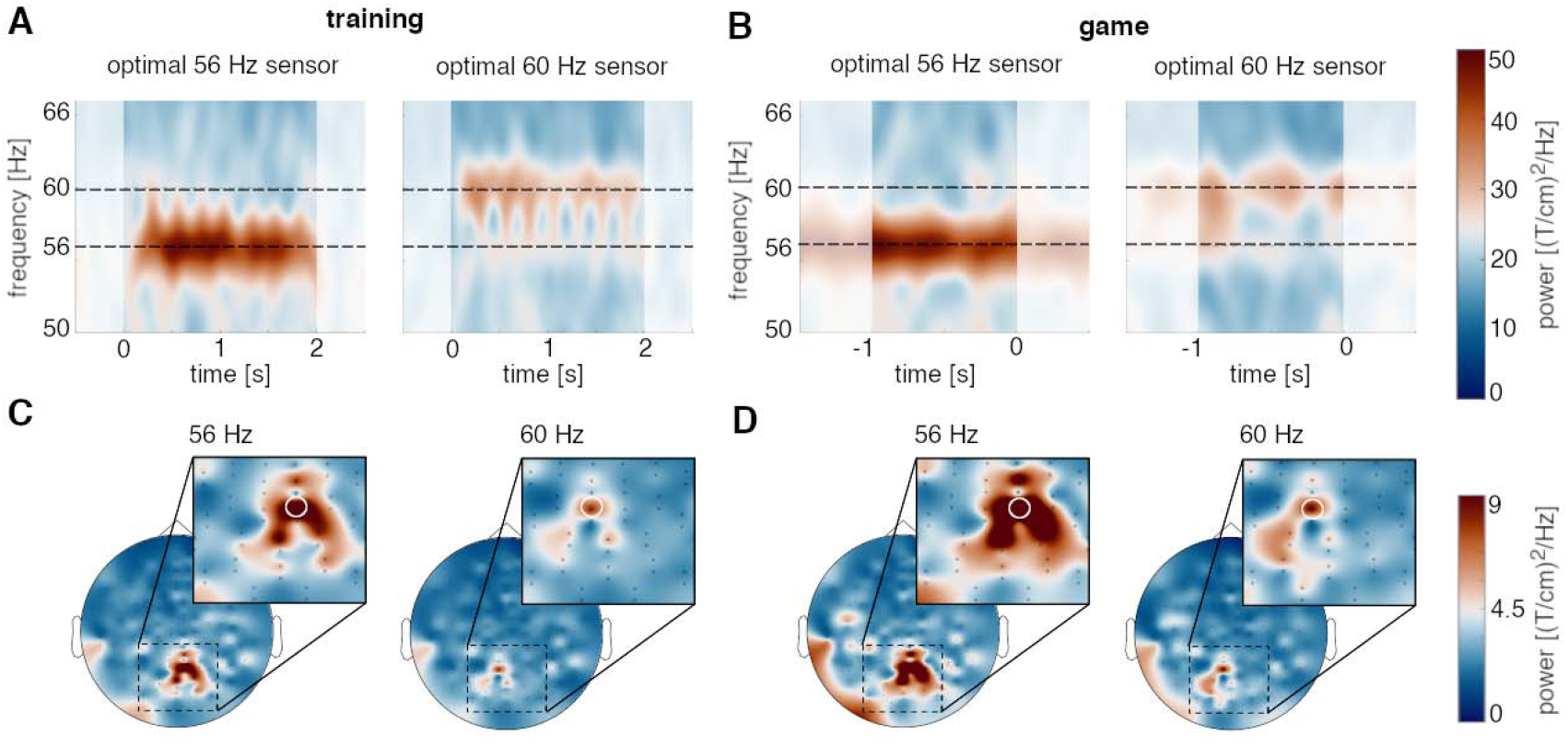
Time-frequency representations and topography of the frequency-tagging response during the training (A, C) and game sessions (B, D). A,B, At the group-level, the gradiometers with the strongest frequency-tagging response showed a clear increase in power at 56 Hz during covert attention-left trials and at 60 Hz during covert attention-right trials. In the training session, time 0 and 2 s denote the on and offset of the frequency-tagging, respectively. In the gaming session, time 0 s denotes the moment of any hit or miss events, while frequency-tagging was uninterrupted. C,D, Topographies of 56 Hz power during attention-left trials and 60 Hz power during attention-right trials revealed spatially distinct and well-localised responses.

### 2.5. Classification algorithm

The data acquired through the training session were used to train a support vector machine (SVM) classifier with a 4-fold cross-validation (Cawley & Talbot, 2002, Hastie, Tibshirani & Friedman, 2008) to decode the direction of covert attention. Considering that the frequency-tagging response is well localised within the primary visual cortex (Zhigalov & Jensen, 2020), we used only seven occipital gradiometers (Fig. 3A) to quantify the modulation of power by covert attention. The power of the frequency-tagging response was computed at 56 and 60 Hz for each gradiometer, resulting in fourteen features. These features along with labels indicating the direction of attention were used to train a linear SVM classifier, as implemented in MATLAB.

**Fig. 3.**
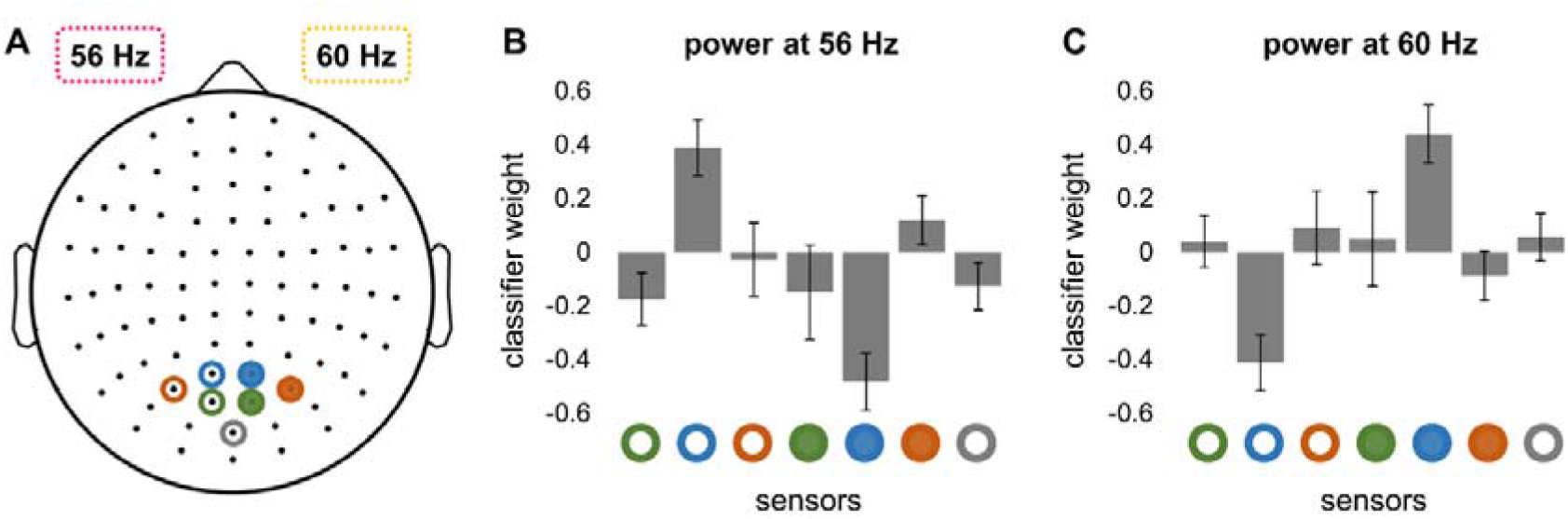
Selected gradiometers and classifier weights at the group level. (A) Seven occipital sensors were used to train the classifier. The patches flickering at 56 and 60 Hz were presented in the left and right hemifields, respectively. (B, C) Classifier weights at the group-level. Bars indicate the standard error of the mean. The three coloured open and filled circles represent left-side and right-side sensors, respectively.

### 2.6. Data processing

Offline processing was performed in MATLAB R2020b using custom-made scripts as well as the Fieldtrip toolbox version 20210106 (Oostenveld et al., 2011). Recordings were kept as raw as possible to reflect the real-time purpose, which included not performing common artifact removal techniques or baseline corrections. Accordingly, the MEG data was divided into epochs and filtered between 1 and 80 Hz. Additionally, a 50 Hz notch filter was applied. Epochs constituted of a range from 1 s before and after frequency-tagging for the training session and 3 s pre to 1 s post pong ball collision events during gaming.

Time-frequency decomposition was performed between 40 and 70 Hz using a Hanning tapered dynamic sliding window covering 20 cycles per frequency in steps of 25 ms. Relevant sensors were identified using an occipital-to-parietal region of interest and subtracting left-trial from right-trial data. For the game data, only successful trials were considered. Perceptually uniform and universally readable colormaps were applied to all visualisations (Crameri et al., 2020).

The information transfer rate was calculated with the following formula (Wolpaw et al., 2002):

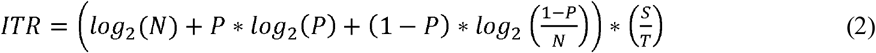

Where *N* is the number of targets, *P* denotes the classification accuracy, *S* is the number of trials and *T* is the time in minutes.

To exploratively estimate any trends between frequency-tagging responses and training as well as gaming accuracy, we performed correlation analyses using Spearman correlation. However, the outcome of the between-subject analysis was not assessed statistically as this would require larger sample sizes.

Lastly, power at 56 and 60 Hz as well as eye-movement data were averaged over a 1 s sliding window in steps of 1 ms to reflect the real-time setup used in the study. The data was then downsampled to 100 Hz and split into right and left trials. Again, for the game session, only successful trials were considered. For both training and game sessions, attention-left and attention-right trials were tested against each other using cluster permutation on dependent sample t-tests for each time point. Right- and left side conditions were shuffled for 1000 iterations to calculate the 5% highest t-sums of randomly formed clusters. Only clusters whose t-sum exceeded 5% of the previously found clusters with the highest t-sum were considered significant.

## 3. Results

In this study, we utilised a novel rapid, invisible frequency-tagging technique in a brain-computer interface paradigm. In a training session, participants were required to covertly attend either the right or the left side of the screen. Their SSVEF responses over seven occipital sensors were used to train a support vector machine (SVM) to distinguish between left and right trials. Afterwards, participants used convert attention to control the position of the bar in the BCI-pong game, testing the efficacy of the trained classifier.

### 3.1. Time-frequency analysis of the frequency-tagging response

We first performed a time-frequency analysis of power of MEG signals, investigating whether the short 1-second timeframe was sufficient to wilfully produce changes in neuronal activity.

During training, participants were instructed to focus their attention to the left or right side of the screen for 2 s while rapid-frequency tagging was applied. At the group level, the power at the gradiometer sensor with the strongest frequency-tagging response clearly increased during the tagging interval at both 56 and 60 Hz and dropped to a baseline after cessation of the tagging (see Fig. 2A). A similar increase in power was observed during the game, where continuous frequency-tagging was applied (see Fig. 2B).

### 3.2. Topography of the frequency tagging response

To assess the spatial distribution of the frequency-tagging response, we computed the spectral power at 56 Hz during covert attention-left trials and at 60 Hz during covert attention-right trials for each gradiometer. It should be noted that the spectral power was assessed during a 2 s tagging interval in the training session, and during a 1 s interval during the game (1 s before the ball collided with the bar or not). At the group level, the response was mainly localised over occipital areas, and the topographies were strikingly similar in training and game sessions (Fig. 2). This suggests that covert attention modulates the neuronal response in the visual cortex locally.

Together, the results indicated that the covert attention robustly modulates the power of frequency-tagging responses within a 1-s interval, meeting the requirements for the BCI-pong game.

### 3.3. Classification and performance

We implemented a linear SVM classifier to decode the direction of covert attention based on the spectral power of the frequency-tagging response. In line with the topography of the strongest responses (see, Fig. 2), only seven occipital gradiometers (Fig. 3A) were selected to train the classifier. The weights of the classifier for 56 and 60 Hz (Fig. 3B,C) showed a roughly inverse pattern (or inverse contribution) as expected for bilateral visual stimulation. Interestingly, the largest weights associated with two bilateral sensors (Fig. 3, blue circles) were highly consistent across participants, suggesting that only two sensors might be sufficient to provide a reasonable classification accuracy at the group level.

The cross-validation accuracy during the training session was above chance level (50%) for seven out of eight participants, and on average it was slightly above 60% (Fig. 4A). The spatially distinct and well-localised responses were again visible when looking at individual participants. Interestingly, the difference in spectral power at 56 Hz (as well as at 60 Hz) between attention-left and attention-right trials at the sensor with the strongest tagging response during training, did not show a clear linear relationship with classification accuracy (Fig. 4B), suggesting that the single gradiometer does not provide sufficient accuracy.

**Fig. 4.**
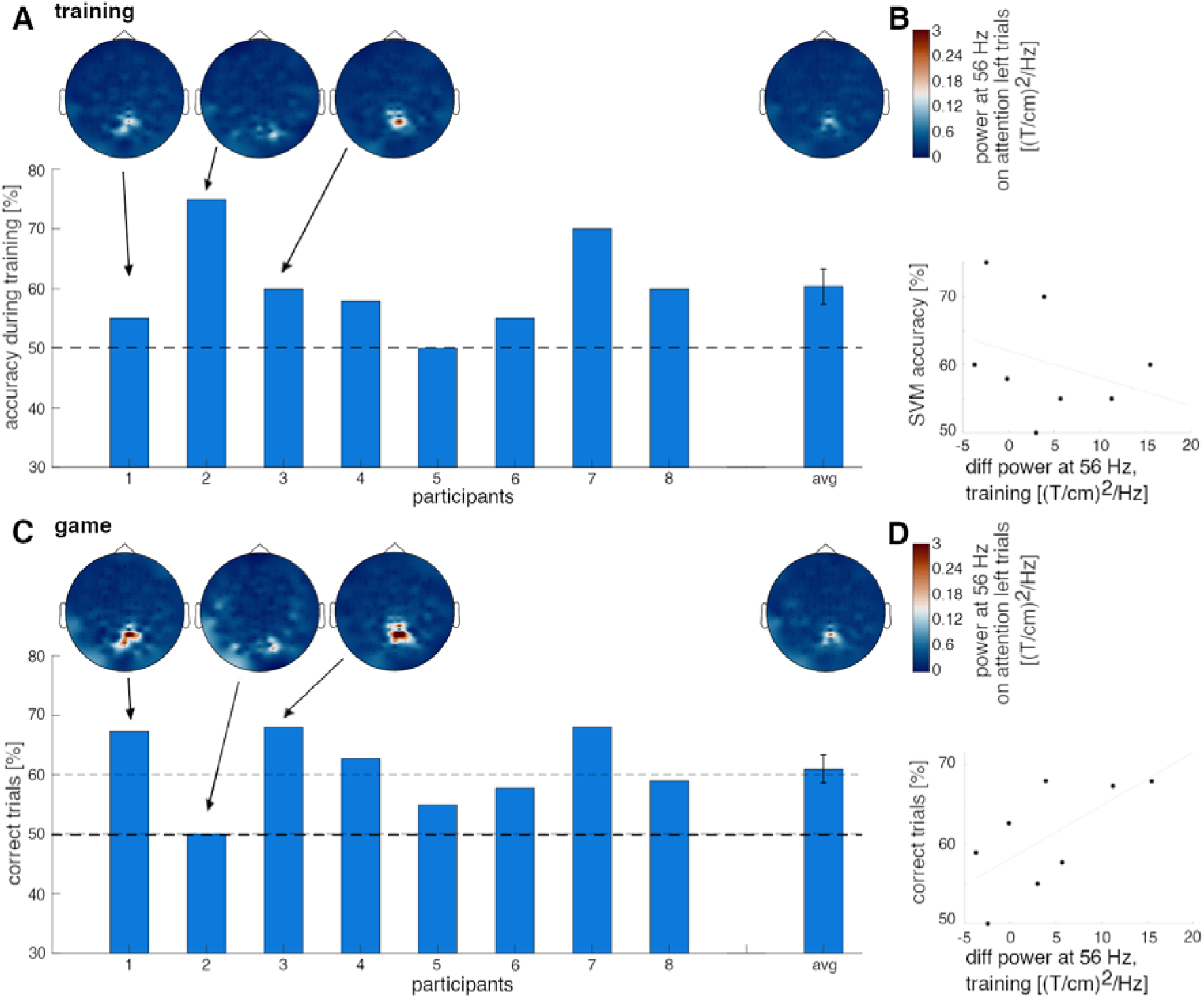
BCI-pong performance. (A) Cross-validation accuracy of the classifier for individual participants and the group average. Topographies of spectral power at 56 Hz are displayed for three representative participants and for the group average. (B) Relationship between attention left versus right power differences at 56 Hz and classifier accuracy. Each dot represents one participant. No clear connection is apparent. (C) Accuracy during the game for individual participants and the group average. Topographies of spectral power at 56 Hz are displayed for three representative participants and for the group average. (D) Performance in the game seems to be related to attention left versus right power differences at 56 Hz during training. Each dot represents one participant.

In real-time game session, 7 out of 8 participants performed above chance level (Fig. 4C). Notably, most participants showed higher accuracy during the game compared to the accuracy estimated based on the training data. As such, the average accuracy exceeded 60%. And 4 out of 8 participants reached accuracies clearly over 70%. Again, topographies showed the same distinct and local pattern as before for individual participants, with stronger power compared to the training session. On average, the information transfer rate of 3.9 bpm was achieved during the game, with a maximum value of 9.8 bpm.

Offline analysis of the MEG data showed that the topography of the frequency-tagging response did not change during the game, suggesting that covert attention modulated neuronal activity mainly in occipital-parietal areas. Interestingly, the difference in spectral power at 56 Hz between attention-left and attention-right training trials at the sensor with strongest tagging response during training, seemed to be highly predictive for the success in the PONG game (r = 0.55, Pearson correlation; Fig. 4D). This might suggest that attentional modulation of spectral power becomes more spatially local during the game.

Overall, our results showed that most participants were able to control covert attention and hence, successfully play the BCI-pong game.

### 3.4 The influence of saccades on BCI performance and frequency-tagging response

To confirm that our BCI in fact relied on covert attention and did not simply reflect amplified frequency-tagging responses due to eye movements towards the flickering targets, we analysed the eye tracker data. During training, there was no apparent difference in saccadic behaviour between left and right trials (Fig. 5A). Yet the spectral power at 56 Hz was higher during attention-left compared to attention-right trials, and as expected, the power at 60 Hz was higher during attention-right trials compared to attention-left trials (see Fig. 5A). We conclude that this difference in the frequency tagging response is explained by covert attention rather than eye movements. The same was true for game sessions. Again, differences in eye movements between right and left trials were not observable (Fig. 5B). The differences in spectral power were significantly enhanced in the gaming session compared to the training session. As such, significant condition differences (p < 0.01, permutation test) between attention-left and attention-right trials were observable in both frequencies spanning from -700 ms prior to and 100 ms after hit or miss events.

**Fig. 5.**
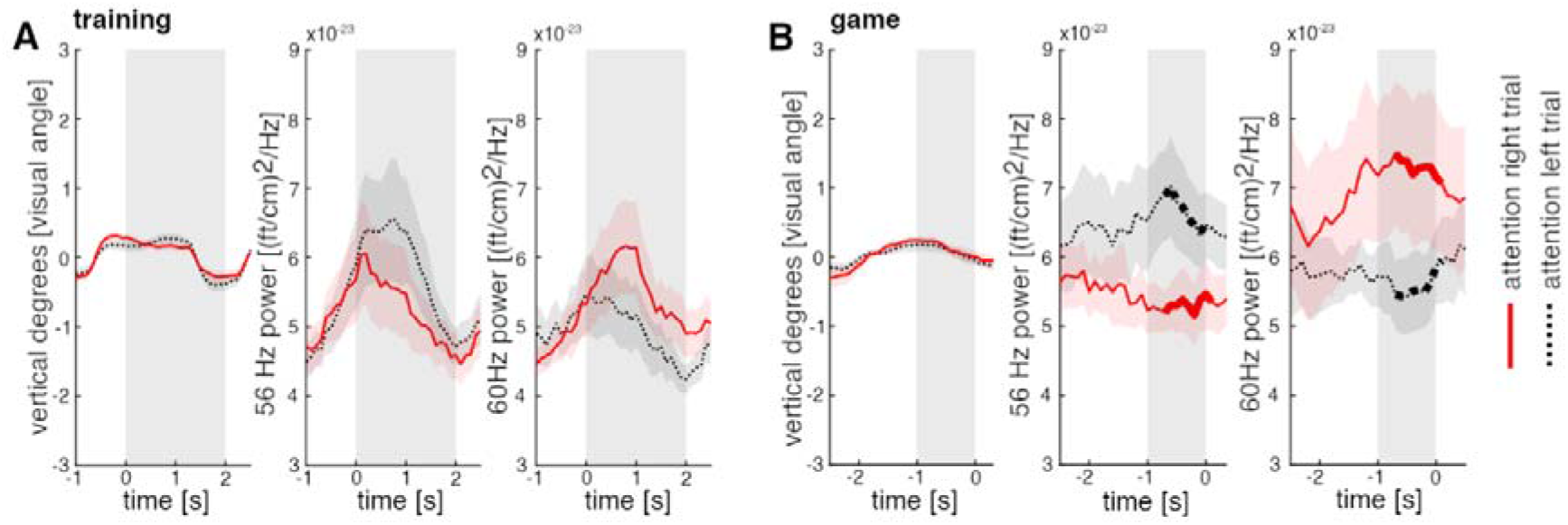
Influence of saccades on BCI -performance during fixation. (A) Changes in eye position and spectral power at 56 and 60 Hz during training. No difference between attention-left and attention-right trials is apparent for eye positions, contrary to the spectral power. Red solid lines indicate attention-right trials, while black dashed lines indicate attention-left trials. Grey areas mark the time interval where frequency-tagging was applied (B) The game session shows no differences in eye positions for attention-left and attention-right trials; however, it reveals larger differences in spectral power for these trials. Significant condition effects are marked in bold. Grey areas mark the time-window used to estimate direction of covert attention.

Furthermore, the eye positions of all participants were relatively stable (Fig. 6A) during the critical period (i.e., the last second before hit or miss) for each trial of the game. The majority of horizontal eye movements remained within 1 degree of visual angle from the fixation cross (on average 97% of eye positions over participants). Furthermore, barely any eye positions were recorded outside of 2 degrees left and right to the fixation cross (on average 1% per participant). Within these 2 degrees, there is no clear pattern of an increase in spectral power to either direction. Aggregating eye positions into 40 bins and averaging the power of the frequency-tagging response does not show a clear pattern nor a difference between 56 Hz and 60 Hz (Fig. 6B).

**Fig. 6.**
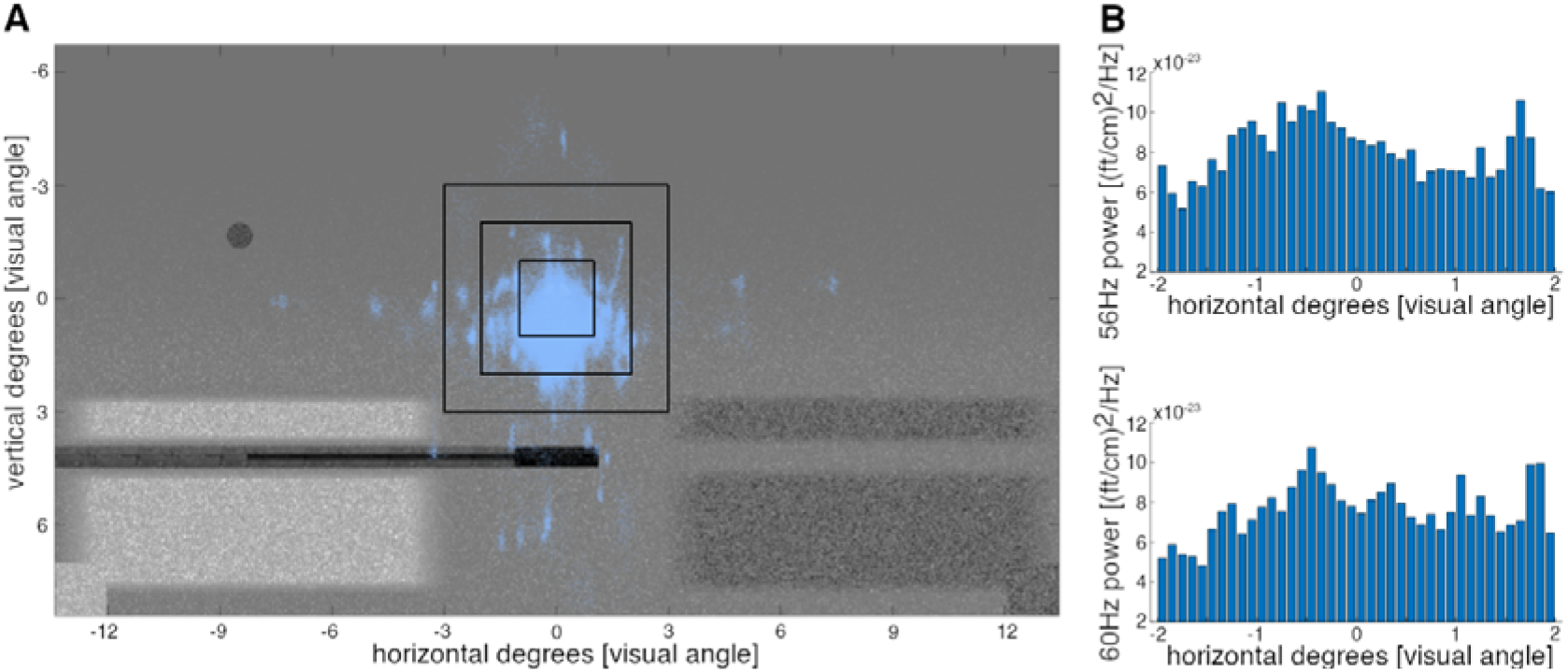
(A) Illustration of the eye positions of all participants across trials during the last second before hit or miss during the game. The black squares indicate 3, 2, and 1 degree of visual angle around the fixation cross, respectively. Especially for horizontal eye movements, most data points stay within 1 degree of visual angle. (B) Histograms illustrating the mean power at 56 Hz and 60 Hz covering 40 binned positions over horizontal eye movements within 2 degrees of visual angle around the fixation cross. No clear association between power of the response and eye-position is observable.

Our data shows that when eye-movement is restricted, it does not affect frequency-tagging responses.

## 4. Discussion

In this study, we have shown that rapid invisible frequency-tagging (RIFT) can be used to decode the direction of covert attention through power estimation of neuronal responses. Our results demonstrate that after a brief training, most participants were able to play the BCI-pong game successfully.

The novel RIFT technique provides several advantages compared to conventional SSVEP/SSVEF based BCI systems (Wolpaw et al., 2002). Here, we highlight three key advantages: invisibility of the flicker, faster detection of neuronal changes, and high signal-to-noise ratio.

Particularly, RIFT produces high-frequency visual flicker that is not perceived by the participant. Accordingly, there is less distraction and strain compared to low-frequency flicker techniques. This is important for both practical applications and fundamental research (Kaplan et al., 2013), as it allows for longer BCI-use by avoiding fatigue related to visual flicker. Additionally, RIFT evokes a robust neuronal response that can be decoded within a short period of time. Our study shows that the high-frequency tagging response modulated by covert attention can be reliably detected within a less-than-one-second window. This property alone can significantly increase the information transfer rate in the existing BCI systems. As such, our system produced higher information transfer rates compared to other covert-attention BCI systems based on alpha activity (e.g., Horschig et al., 2014 achieved an average of 2 bpm compared to 3.9 bpm in this study). Although, in this study the analysis window was limited by one second, there is a potential to further reduce the window, while preserving the decodability of responses. Finally, high-frequency flicker evokes a neuronal response with high signal-to-noise ratio since the response is not affected by the pronounced endogenous rhythms at low frequencies. In contrast, when using low-frequency visual stimulation, there is a chance that the stimulation frequency may be in a close proximity to endogenous rhythms (Haegens & Zion Golumbic, 2018; Jensen & Mazaheri, 2010), which could lead to attenuation of and interference with the neuronal response (Keitel et al., 2014; Spaak et al., 2014).

Eye-movements during BCI control might raise concern as to whether saccades towards the frequency-tagging stimuli rather than covert attention modulated the neural responses. We showed that while attention left- and right-trials significantly differed in power, the saccadic eye movements were indistinguishable in these trails. This is important as it emphasises that covert attention could not simply be reduced to eye-movements towards the frequency tagged objects. On a smaller scale, directional biases in microsaccades are correlated with neural signatures of covert spatial attention (Lowet, 2018, Neuron). However, a recent study revealed that this link was not causal in monkeys (Ya et al., 2022).

Considering a future application of RIFT BCI for highly mobile BCI applications, we specifically designed our BCI system using only a small number of MEG sensors and with minimal processing requirements. As we could show that the RIFT response is strongly localised, it can be captured by only few sensors. Although, in this study we used seven sensors to control the BCI, the group level analysis of classification weights suggested that already two bilateral sensors in the occipital region provide a reasonable classification accuracy. This shows that our BCI can be easily implemented in mobile and flexible systems for example with novel optically pumped magnetometers (OPMs), which eases application for children and clinical populations, while being less restrictive on movement compared to conventional measuring systems.

### Limitations of the study

First, for this proof of principle, we recruited a limited number of participants (N=8) and implemented short training sessions. This can be easily expanded to a larger cohort of participants enabling more rigorous statistical analysis.

Second, we used a simple decoding technique, linear SVM. More sophisticated machine learning approaches may provide a better classification accuracy and hence, improve the performance in the game.

Third, we used a traditional MEG system to record the brain activity, which due to the rigid position of the sensors, may provide slightly inconsistent results related to the participant’s head movement during the experiment. Moreover, such MEG systems have poor mobility, and hence, the practical application may be limited. Recent development in OPM shows that magnetic fields of the brain can be recorded using highly flexible OPM sensors (Boto et al., 2018; Brooks et al., 2021), and thus, the problem of rigid sensors position, and poor mobility can easily be overcome.

### Future work

The use of rapid invisible frequency-tagging has strong advantages compared to existing BCI systems. However, future work can further improve the application of this technique.

Our study shows that covert attention modulates frequency-tagging responses locally, and thus, a very limited number of sensors is necessary to detect the neuronal responses of RIFT. In this case, OPM sensors provide an excellent solution, since they do not constraint the participant movement, and importantly, such sensors can be placed flexibly and have a much lower cost compared to conventional MEG systems.

## 5. Conclusion

This study provides novel insights into brain-computer interfaces based on covert attention. We applied a new rapid-frequency tagging technique that has two key advantages compared to the traditional methods. First, it does not produce visible flicker and hence less distractive to participants. Second, it allows decoding attentional modulation of the neuronal response within a short period of time and hence, enables high-speed communication through increased information transfer rates. Combining the RIFT technique with novel wearable OPM sensors will allow making this BCI system more mobile and move development of BCIs to the next level.

## Acknowledgements

The work was supported by the following funding: a James S. McDonnell Foundation Understanding Human Cognition Collaborative Award (grant number 220020448), the Wellcome Trust Investigator Award in Science (grant number 207550), a BBSRC grant (BB/R018723/1) as well as the Royal Society Wolfson Research Merit Award.

## Competing interests

There are no competing interests

## Notes

### Competing Interest Statement

The authors have declared no competing interest.

